# Content-Based Similarity Search in Large-Scale DNA Data Storage Systems

**DOI:** 10.1101/2020.05.25.115477

**Authors:** Callista Bee, Yuan-Jyue Chen, David Ward, Xiaomeng Liu, Georg Seelig, Karin Strauss, Luis Ceze

## Abstract

Synthetic DNA has the potential to store the world’s continuously growing amount of data in an extremely dense and durable medium. Current proposals for DNA-based digital storage systems include the ability to retrieve individual files by their unique identifier, but not by their content. Here, we demonstrate content-based retrieval from a DNA database by learning a mapping from images to DNA sequences such that an encoded query image will retrieve visually similar images from the database via DNA hybridization. We encoded and synthesized a database of 1.6 million images and queried it with a variety of images, showing that each query retrieves a sample of the database containing visually similar images are retrieved at a rate much greater than chance. We compare our results with several algorithms for similarity search in electronic systems, and demonstrate that our molecular approach is competitive with state-of-the-art electronics.

**One Sentence Summary:** Learned encodings enable content-based image similarity search from a database of 1.6 million images encoded in synthetic DNA.

## Main Text

Interacting with digital documents is a staple of modern society. Since the dawn of online social media, the amount of text, images, and videos that the world produces has been growing rapidly in an exponential fashion, outpacing the capacity growth of traditional storage media (solid state, magnetic, optical). To address this gap and store this ever-expanding library of data for future generations, researchers are developing practical systems using synthetic DNA as a high density and durable storage medium (*1–7*), with several orders of magnitude higher density and durability than current storage technologies. An important feature of these systems is the ability to retrieve single documents without having to sequence a large, dense pool of data in molecular form. To accomplish this, these systems leverage the specificity of DNA hybridization to implement key-based retrieval, where a short, predetermined sequence is used to retrieve the document associated with that sequence (*5–6*).

Although key-based retrieval might be sufficient for a well-maintained library or archive, modern search and recommendation systems do not assume users know the exact key of the document they are looking for, and thus make heavy use of content-based retrieval. Moreover, similarity search is an important component of modern machine learning pipelines. Early formulations of DNA databases (*8–11*) proposed that hybridization could also be used to search through the content of the documents in the database. However, robust and efficient encoding of arbitrary digital data in DNA requires randomizing the document content (*12*), which impedes direct content-based search. Electronic database systems separate the concerns of search and storage by maintaining an index structure that is designed to facilitate content-based retrieval. We follow this design principle by creating a DNA-based index that is optimized for *similarity search*, where an example document is used to retrieve similar documents from a database. Augmenting the density and longevity of DNA data storage with the computational power of similarity search will enable applications that are infeasible for electronic systems. For instance, an easily-searchable archive of today’s primary source material would be invaluable to future historians seeking to explore this critical stage of human history. While it would be require too much power, space, and bandwidth to build electronically, a DNA-based archive could be replicated and distributed worldwide with minimal resources.

Document similarity is typically formulated as a geometric problem (*13*) where each document is converted to a vector in a high-dimensional *feature space*, with the property that neighboring feature vectors represent subjectively similar documents. This conversion process is called *feature extraction*, and there are a variety of methods that work well, depending on the type of document and the goals of the application. For documents like images, intermediate layer activations from neural networks trained on image classification problems (such as VGG16 (*14*) or AlexNet (*15*)) tend to perform well as feature vectors for similarity search tasks (*16*). Figure 1A illustrates this with a two-dimensional *t*-SNE (t-distributed Stochastic Neighbor Embedding) projection of 4096-dimensional feature vectors extracted using the FC2 layer of VGG16.

**Fig. 1.**
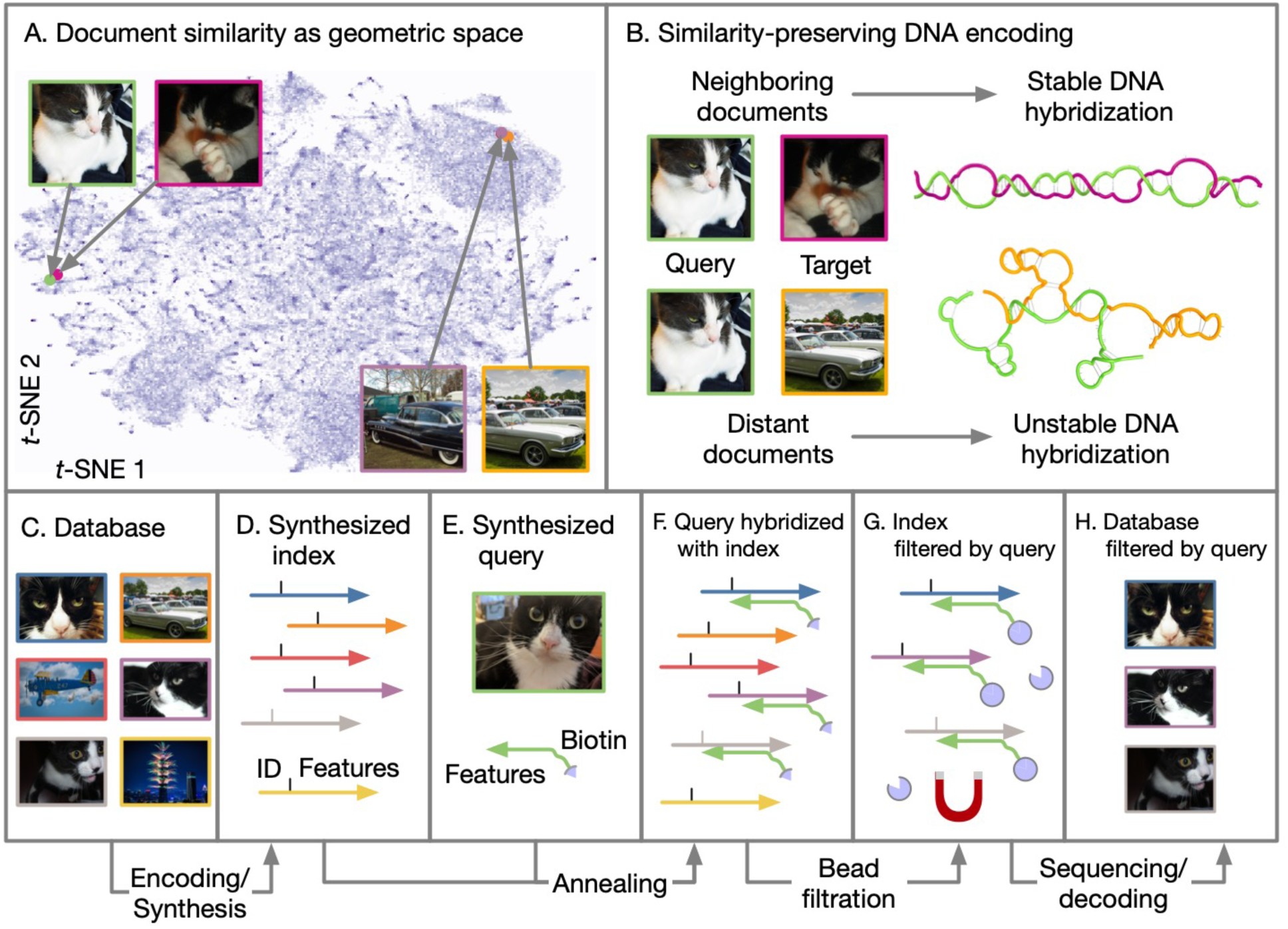
Overview DNA-based similarity search. (A) Illustration of a feature space where neighboring documents are subjectively similar. (B) A similarity-preserving DNA encoding is one where the reverse complement of a query document’s sequence hybridizes with a neighboring target document’s sequence, but not with a distant target’s. Note that the query is color-coded green and the targets with other colors. (C-H) The retrieval process. A database (C) is encoded and synthesized to produce an DNA-based index (D). Arrowheads indicate 3’ ends of DNA. An encoded query (E) is annealed with a sample of the database (F), which is filtered with magnetic beads (G). The filtered database is sequenced to reveal the IDs of retrieved documents, which are used to look them up in the original database (H).

Searching through such a high-dimensional space is challenging for electronic systems. The “curse of dimensionality” (*13*) means that exact indexing schemes in high-dimensionality spaces are no better than a costly linear or “brute force” search, which is infeasible for large databases. Instead, efficient similarity search algorithms perform an approximate search that allows for errors in the results. We adopt that relaxation as well. Rather than finding exact nearest neighbors, our goal is to maximize the number of near neighbors retrieved while minimizing the number of irrelevant results.

To accomplish this, we want to encode feature vectors as DNA sequences such that single stranded molecules created from an encoded target and the reverse-complement of an encoded query are likely to form stable hybridized structures when the query and target feature vectors are neighboring, but not when they are distant (Figure 1B). Given such an encoding, we can construct an index where each document is associated with a single strand of DNA that contains the document’s ID, alongside with its encoded feature vector (Figure 1C-D). An encoded query (Figure 1E) can then be used as a hybridization probe to filter out similar documents from the index (Figure 1F-G). The filtered index can be sequenced and decoded to recover the IDs of documents that are similar to the query. These documents can then be retrieved from a key-based database and displayed to the user (Figure 1H).

Designing a universal encoding from feature vectors to DNA sequences is difficult because of the high dimensionality of both feature space and DNA sequence space, and because of the non-linear nature of DNA hybridization. An alternative is to use machine learning to optimize the encoding for a particular dataset. Early researchers (*9*) achieved this by clustering the dataset using *k*-means, then mapping each cluster center to a known DNA sequence which is assigned to each document in that cluster. By reducing content-based retrieval to exact key-based retrieval, their approach sidesteps any issues with unwanted DNA hybridization. However, once the cluster labels are chosen, they are static; any additional data added to the database must fit into the existing clusters, even if it would cause the cluster centers to change.

Rather than using a codebook of pre-determined sequences, our technique leverages neural networks to map feature vectors directly onto fixed-length DNA sequences. A key component of our technique is a differentiable model of DNA hybridization that is trained alongside the encoder. This allows us to optimize the encoder to minimize unwanted hybridization while preserving the flexibility of inexact matching. To train the predictor, the reactions between pairs of sequences output by the encoder are simulated via NUPACK (*17*). We consider this simulation to be the “gold standard” for computing the outcome of a reaction, but it cannot be used directly to train the encoder, because its computation is not differentiable.

We chose to focus first on encoding feature vectors derived from images, as similarity between images is easy to visualize, and large datasets are readily available. We use OpenImages (*18*), a collection of roughly 9 million images. Figure 2A illustrates the neural network that maps images to DNA sequences. First, we use the FC2 intermediate layer of VGG16 (*14*) to extract a feature vector. A fully-connected neural network with a single hidden layer converts these feature vectors to “one-hot” DNA sequences, where each position is represented numerically by a four-channel vector (one channel for each possible base) whose components sum to one. More details on the encoder architecture are available in the supplementary material.

**Fig. 2.**
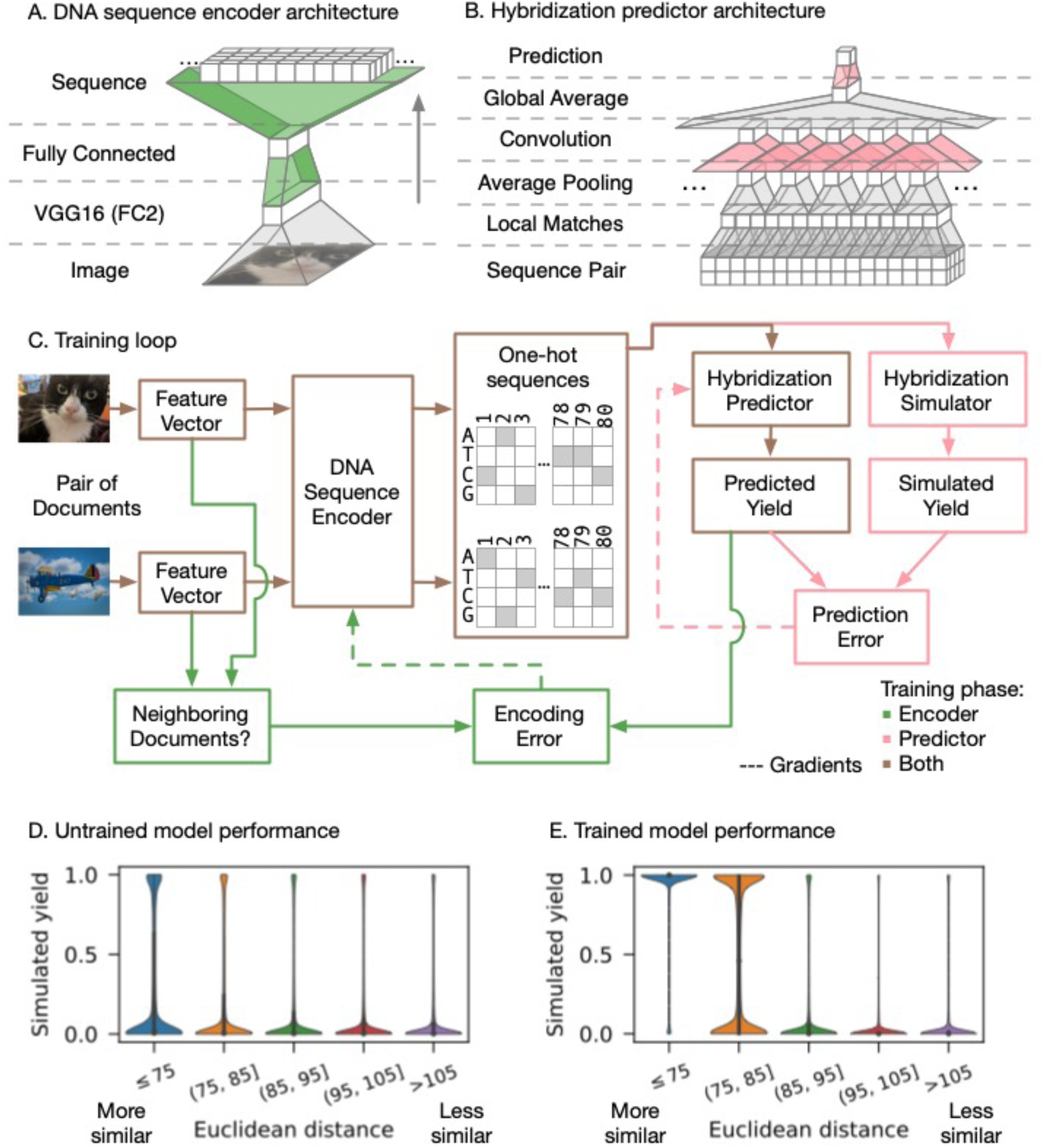
Overview of our neural network architectures and training process. (A-B) The neural network architectures for the image-to-sequence encoder, and the hybridization predictor. Boxes represent layers, and trapezoids represent operations to go from one layer to the next. Only the colored operations have parameters that change during training. (C) The training loop for the neural networks. Lines indicate data flow; dashed lines indicate parameter gradients calculated using backpropagation. Green indicates operations only performed during encoder training, while pink indicates operations used only during yield predictor training. All other operations are used in both training phases. (D) Simulated performance of an untrained model, evaluated on 1.6 million random pairs of images. Each violin depicts the distribution of simulated hybridizations for pairs whose feature vectors’ Euclidean distance lie within a certain range. (E) Simulated performance of a trained model, evaluated on the same set of random pairs.

The hybridization predictor (Fig. 2B) takes a pair of one-hot sequences and predicts the yield of the hybridization reaction between the top sequence and the reverse-complement of the bottom sequence. It first pre-processes the sequence pair with a sliding window operation that extracts local matches (Fig. S3B). Following local average pooling, a trainable convolutional layer enables the predictor to weigh different kinds of local matches differently. Because we are not predicting where hybridization occurs, a global average pooling operation removes spatial relevance. The prediction is then computed with a logistic regression (in our experience, this is more effective than a linear regression because most hybridization yields are either close to 0 or close to 1). More details on the yield predictor architecture are available in the supplementary material.

Figure 2C outlines the training procedure, which alternates between encoder and predictor training phases. In the encoder training phase, a pair of image feature vectors are compared to determine if they are close enough to be deemed “similar”. The threshold for similarity is determined based on subjective evaluation of the feature space (see Figure S1 in the supplementary material). The feature vectors are encoded independently to produce a pair of one-hot sequences, which are passed to the hybridization predictor. If the predicted yield is low and the documents are similar, or the predicted yield is high and the documents are not similar, the encoder’s parameters are modified (via gradient descent) to correct the error.

In the predictor training phase, the one-hot sequences output by the encoder are discretized and their reaction is simulated with NUPACK. If the predicted yield differs from NUPACK’s simulation result, the predictor’s parameters are modified (via gradient descent) to correct the error. Eventually, the predictor learns a model of hybridization that is good enough for the encoder to learn how to map feature vectors to DNA sequences, such that neighboring feature vectors are much more likely to result in pairs of sequences that hybridize (Figure 2D-E).

During training, we withheld a fixed subset of 1.6 million images from OpenImages V4 to be used as our “database” for laboratory experiments. After training our encoder, we transformed each of these images into a DNA sequence using the trained encoder. In addition to the encoded features, each image’s sequence contains a unique, decodable barcode that can identify the sequence without using reference-based alignment, as well as conserved regions to facilitate amplification and processing via PCR (polymerase chain reaction). Each image’s sequence is short enough to fit on a single synthesizable DNA oligomer (see Figure S4 in the supplementary material).

Our query images did not come from the OpenImages dataset. To conduct similarity search with an image, we ordered a biotinylated probe oligomer that contains the reverse complement of the query’s encoded feature sequence. We annealed the probe with a sample of the database, and then separated the annealed target/query pairs from the database using streptavidin-conjugated magnetic beads. We then use high-throughput sequencing to reveal which database sequences persist in the separated mixture, and measure how frequently each of them occur.

Figure 3 shows the experimental results for three different query images. If we consider images with sequencing read counts above a certain threshold to be “retrieved”, we can characterize the set of retrieved images for a variety of thresholds. Figure 3A shows that higher read counts are associated with sets of images that are closer to the query in Euclidean distance. We can quantitatively characterize the quality of a retrieved set by its recall of the 100 nearest neighbors; that is, the number of images in the set that are among the 100 most similar images to the query in the database. Figure 3B shows that, as the read threshold increases, the number of total images in the retrieved set drops very low before you begin to sacrifice nearest neighbor recall. We can also visually inspect the retrieved set by sorting its contents and displaying the most similar images. Figure 3C shows that, even with very aggressive filtering, the retrieved set still contains images that are relevant to the query. If the read counts for each image are proportional to their concentrations in the filtered mixture, this means that the filtered mixture could be diluted about 1000x, conserving sequencing resources while still retrieving relevant images.

**Fig. 3.**
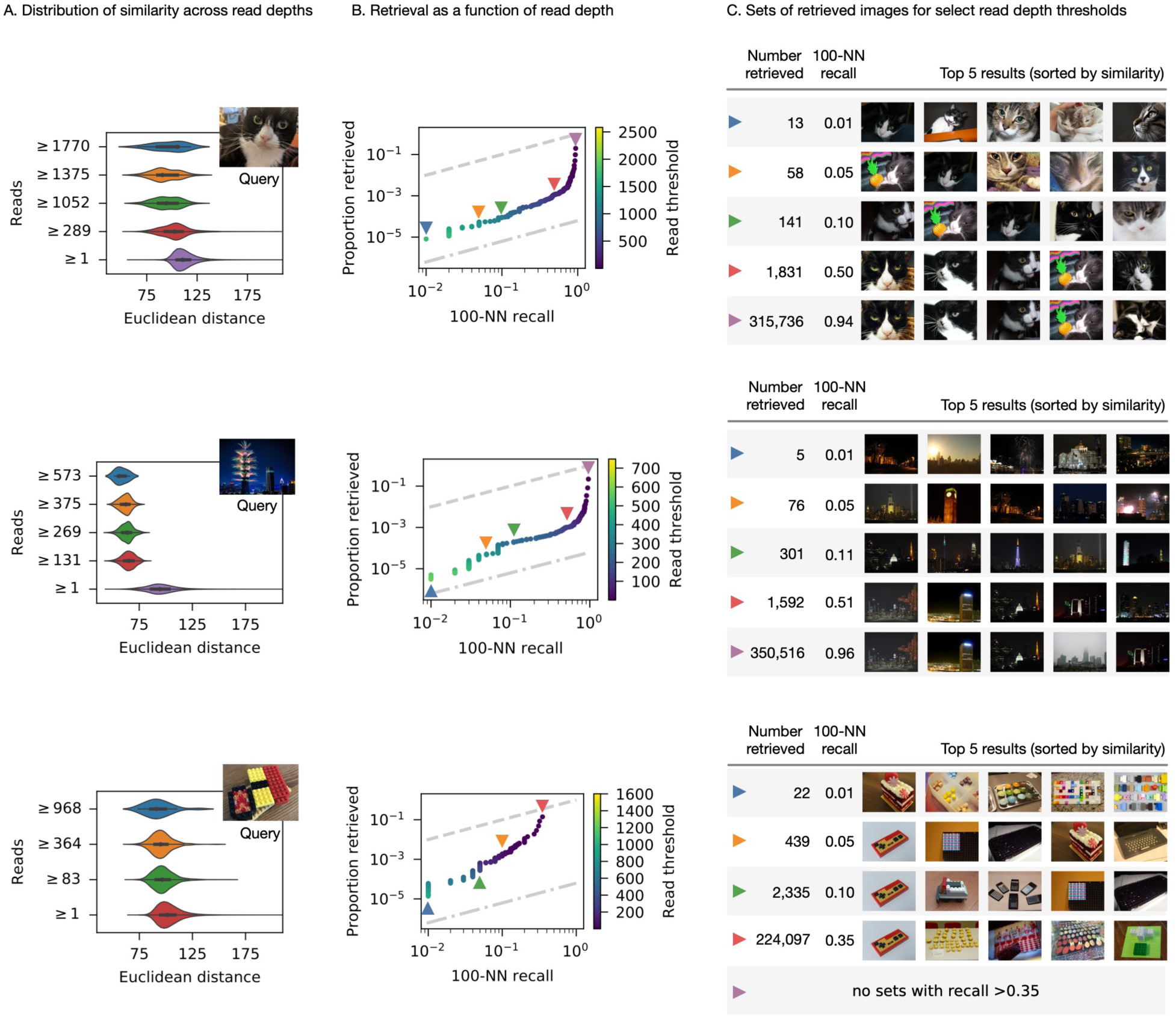
Experimental results for three different query images. Janelle, the cat (top), a building with fireworks (center) and Lego pieces assembled in the shape of sushi (bottom). (A) Distribution of Euclidean distances to the query image, among sets of images with sequencing read depth above a certain threshold. (B) The proportion of the entire dataset that must be retrieved (y-axis) to retrieve a certain proportion of the 100 most similar images (x-axis). Each point represents a threshold for which images with read depth above that threshold are considered “retrieved”. The dashed line indicates chance performance, while the dashed-and-dotted line indicates perfect performance. Colored triangles indicate the thresholds depicted in the other subfigures. (C) The top 5 closest images to the query from result sets where images above a certain read depth threshold are considered “retrieved”.

The performance of a similarity search algorithm can be summarized by the curve in Figure 3B, which measures the proportion of the database that must be retrieved and sorted to achieve a particular 100-nearest neighbor recall. The dashed line above the curve illustrates a “naive” algorithm that randomly samples the database. To retrieve half of the hundred nearest neighbors, it must retrieve half of the database. The dashed-and-dotted line below the curve illustrates a perfect “oracle” algorithm. To retrieve half of the hundred nearest neighbors, it would retrieve exactly those 50 images from the 1.6 million in the database.

Figure 4 places the curve from Figure 3B in context alongside several state-of-the-art *in silico* algorithms that were benchmarked using the same query and same database for each of the queries we evaluated experimentally. Implementations of HNSW (hierarchical navigable small world) graphs (*19*) are among the top performers on approximate nearest neighbor benchmarks (*20*). HNSW requires building and storing a very large index, which may be difficult to scale to large databases. We also tested a quantized version of HNSW with lower memory utilization, developed by Facebook (*21*) (“faiss”, shown in red), annoy (*22*) (shown in green), a popular algorithm developed by Spotify, and RPForest (*23*) (shown in purple), an algorithm designed for the lowest possible memory utilization. While there is room for improvement, our experimental performance is comparable to the state-of-the-art, indicating that DNA-based similarity search is a viable technique for searching the databases of the future.

**Fig. 4.**
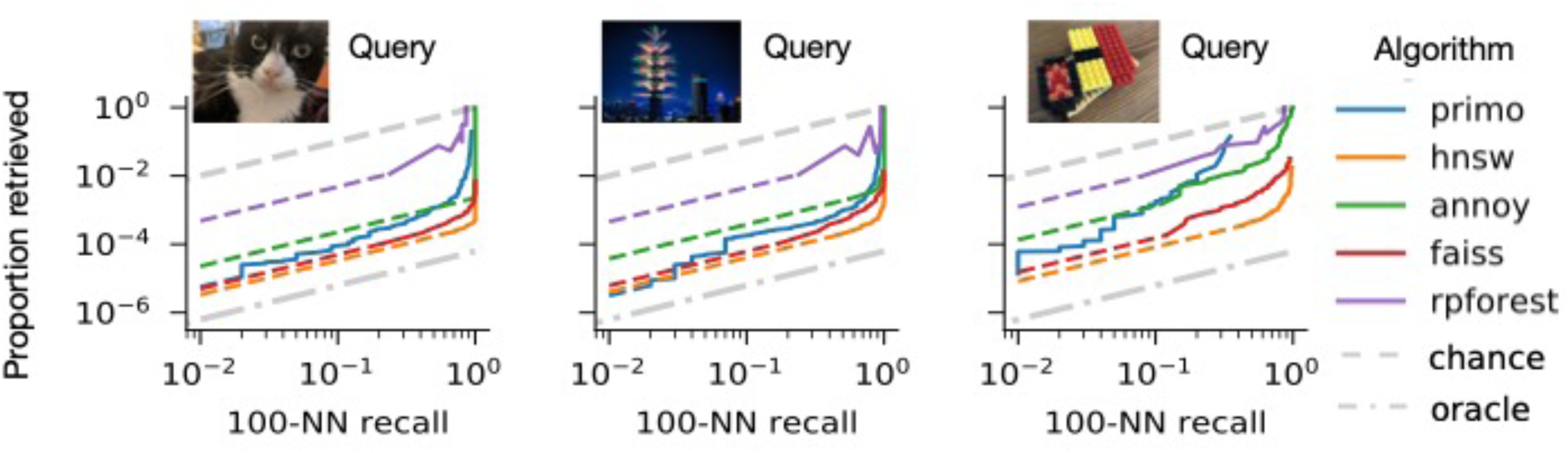
Comparison of our technique (“primo”, shown in blue) with state-of-the-art algorithms for in silico similarity search. Dashed grey and dashed-and-dotted grey lines represent chance performance and perfect performance, respectively. Not all of the algorithms could produce results towards the lower-left (low recall and low proportion retrieved). We assume these algorithms could be stopped early to produce fewer results with a linear decrease in recall; dashed continuations represent these linear interpolations.

This paper detailed the first demonstration of similarity search of digital information in DNA and compared its potential efficiency with electronic systems. The results suggest that, as DNA data storage becomes more practical and scales to larger data sets, similarity search in DNA form is an attractive possibility compared to electronic systems. Combining DNA data storage with similarity search support may offer a path to viable hybrid molecular-electronic computer systems.

## Acknowledgments

The authors thank the Molecular Information Systems Lab for invaluable feedback on the research ideas and results. Dr. Anne Fischer provided several suggestions for data analysis, results presentation and comparisons with electronic systems.

## Author contributions

C.B. developed core learning-based encoding, designed experiments, evaluated results, and wrote manuscript text. Y.C. developed molecular biology protocols, performed experiments and analyzed results. D.W. and X.L. performed experiments. G.S., K.S., and L.C. supervised the work. L.C. suggested initial idea of DNA-based similarity search.

## Data and materials availability

Target images are publicly available via Google: https://storage.googleapis.com/openimages/web/download_v4.html. VGG16 feature vectors can be extracted from images using a publicly available model (via Keras): https://keras.io/api/applications/vgg/#vgg16-function. Code for our trained encoder (to convert feature vectors to DNA sequences) will be available soon. Meanwhile, encoded DNA sequences, query images, sequencing reads, and analysis are available via our repository: https://github.com/uwmisl/primo-data.

## Supplementary Materials

Materials and Methods

Figures S1-S5

## Materials and Methods

### Datasets

With the exception of the query images, all images were collected from Open Images V4, a dataset of over 9 million URLs for images with Creative Commons licenses. Of these, approximately 1.7 million are hosted by the CVDF and available for download; the rest are raw Flickr URLs and may or may not be available. For the image database used in our experiments, we took 1.6 million images from the hosted set. For training, we took images from the full set of 9 million that were not used for training, testing, or experiments.

### Feature Extraction

To extract image features, we processed each image with VGG16, a convolutional neural network designed for image classification. The weights were loaded from the publicly available trained model and left unchanged during our processing. We used the activations of FC2 (the second fully-connected layer) as 4096-dimensional feature vectors.

Figure S1 illustrates the relationship between subjective image similarity and feature vector Euclidean distance. Pairs of images with Euclidean distance of 75 or less tend to be consistently similar, so during training we label these pairs as “similar” and all other pairs as “not similar”.

### Sequence Encoding

The sequence encoder is a fully-connected neural network. Its topology is depicted in Figure S2. The 4096-dimensional FC2 vectors are fed into a 2048-dimensional hidden layer with a rectified linear activation, followed by an output layer with a “one-hot” sequence representation that is 80 nucleotides in length. In this representation, each sequence position has four channels, one for each base. A softmax activation function is applied that forces each position’s channels to sum to 1. A DNA sequence can be read off by picking the channel with the maximum activity at each position.

The one-hot representation can produce indeterminate bases (for example, if all four channels at a position have a value of 0.25). Because of this, a regularization is applied during encoder training to minimize the entropy at each position. This encourages each position to have a well-defined maximum, which improves the accuracy of the yield predictor.

The yield predictor takes a pair of one-hot sequence representations and produces an estimate of the yield of the hybridization reaction between the first sequence and the reverse-complement of the second sequence. It is structured as a convolutional neural network (Figure S3A). The network makes use of a novel local match layer (Figure S3B) that produces vectors of possible matches between each window of 3-mers. This encourages the predictor to make use of any unaligned matches between the two sequences.

### Training

During each round of encoder training, we draw a batch of pairs of feature vectors from the training set where half of the pairs are labeled “similar” (the Euclidean distance between the feature vectors in the pair is 75 or less). The batch of pairs is processed by the encoder, which outputs pairs of one-hot sequences. These are then processed by the yield predictor, which outputs the estimated yield of the hybridization reaction between the first sequence and the reverse complement of the second sequence. The estimated yield of each pair in the batch is used along with the similarity labels (0 for “not similar” and 1 for “similar”) to compute the mean cross-entropy for the batch. We use the cross-entropy as loss function because it penalizes similar images with low estimated yield, dissimilar images with high estimated yield, and any estimated yields that are neither high nor low. The parameters of the encoder are modified (via gradient descent) to minimize the mean cross-entropy. The yield predictor’s parameters are not changed during encoder training.

In order to use gradient descent, the one-hot sequence representations cannot be fully discrete. This can create positions with indeterminate bases, which may interfere with the yield predictor. To discretize the one-hot sequences as much as possible, we add an additional regularization term to the encoder to minimize the per-position entropy of the one-hot sequences. This regularization must be gently applied; if it is too strong, the gradients will explode and training will fail.

During each round of yield predictor training, we draw a random batch of pairs of feature vectors (unlike the encoder batches, these are not conditioned to have a particular distribution of similarity). The batch is processed as above to output the estimated reaction yield.

To simulate the reaction yield with NUPACK, the pairs of sequences are discretized. The first sequence in the pair is treated as the target, and a reverse primer is appended as in Fig. S4A. We do not append the forward primer, barcode, or internal primer as these regions will be double-stranded during retrieval. The second sequence in the pair is treated as the query, and six bases of the reverse primer are appended, and the sequence is reverse-complemented, as in Fig. S4D. The target and query sequences are processed with NUPACK at 21°C using default DNA parameters and an equal molar concentration of 1 nM for both query and target. The simulated reaction yield is computed by dividing the final concentration of the query-target duplex by the initial concentration of 1 nM.

We compute the cross-entropy between NUPACK’s simulated yield and the predictor’s estimated yield for each pair in the batch. The parameters of the yield predictor are modified (via gradient descent) to minimize the mean cross-entropy for the batch. The encoder’s parameters are not changed during predictor training.

### Barcodes

Document IDs are integers in the range 0 to 16,777,215. Because our database only contains 1.6 million entries, we space out their IDs by mapping them to an ID in the full range using a pseudo-random permutation. To construct a DNA barcode from an ID, the randomized ID is first split into to four six-bit symbols. These are encoded with a Reed-Solomon error-correcting code to produce a codeword with six symbols. Each symbol is converted into a five-nucleotide homopolymer-free DNA subsequence using a codebook with 64 entries. The final 30-nucleotide barcode is the concatenation of these subsequences.

To decode a 30-nucleotide barcode, it is split into six five-nucleotide components, and each step of the code is reversed. Limited substitutions can be corrected, but if the sequence cannot be decoded, or it decodes to an ID that is unused, it is rejected as an invalid barcode.

### Oligo Layout

Figure S4 depicts the layouts of our synthesized DNA oligomers, as well as the layouts of double-stranded complexes formed during processing. Each document in the database is associated with a single DNA oligomer (Figure S4A) that contains the barcode and feature regions that are unique to that document. In addition to these unique regions, each database oligo contains three conserved regions (denoted FP, RP, and IP) that are the same across all documents. PCR with FP and RP* is used to create additional copies of all database strands (Figure S4B), to prepare for hybridization and sequencing. Linear PCR with IP* is used to create partially-double stranded copies of each database strand that leave the feature region exposed (Figure S4C).

Figure S4D depicts the layout of a query oligo. During retrieval, the reverse complement of the query document’s features are synthesized along with a 5’ biotin, a short spacer, and the reverse complement of first six bases of RP (which serve as a hybridization toehold). The biotinylated query can react with the exposed feature region of any database oligo (Figure S4E). If the resulting complex is sufficiently stable, it can be filtered from the rest of the database using streptavidin-conjugated magnetic beads.

### Benchmarking of in silico algorithms

The *in-silico* similarity search algorithms we compare against in Fig. 4 all perform an Approximate Nearest-Neighbor (ANN) search. Given a dataset, they create an index structure that is traversed using a given query’s feature vector, to retrieve the documents whose feature vectors are nearest to that query. An approximate search does not scan the entire database, but this may cause it to miss some of the nearest neighbors. We define the *candidate set* as the subset of the database that is scanned (i.e., retrieved from memory and compared to the query). The candidate set is analogous to the set of “retrieved” documents we define by varying the read depth threshold for our lab experiments.

For each algorithm in Fig. 4, and for each of the three queries, we collected the candidate sets for a variety of algorithm-specific parameters that give users control over the specificity of the ANN search. The size of each candidate set (divided by the size of the full database) gives us the proportion retrieved (the y-axis of Fig. 4), whereas the number of true nearest neighbors in each candidate set (out of 100) gives us the 100-nearest-neighbor recall (the x-axis of Fig. 4).

Some algorithms could not retrieve candidate sets below a certain size for any of the attempted parameters. For these, we assume that uniform subsampling would equally limit both the size of the candidate set and the number of nearest neighbors retrieved. This assumption gives us the dashed colored lines for each algorithm in Fig. 4.

### Reagents

The 1.6 million oligos that make up the database were ordered from Twist Bioscience. Biotinylated probe oligos were ordered from IDT. Streptavidin-coated magnetic beads (Dynabeads MyOne Streptavidin T1) was purchased from Thermo Fisher Scientific. USER enzyme was ordered from New England Lab.

### Laboratory Protocol

The general workflow of a similarity search experiment is divided into 8 steps (Figure S7): (1) enrichment of a synthesized oligo pool using PCR, (2) linear amplification of the pool using a forward primer, (3) linear amplification using an internal primer, (4) hybridization experiment using a query strand, (5) magnetic bead extraction, (6) releasing of bead captured strands using digestion of USER enzyme, (7) PCR enrichment of the released oligos (8) ligation to Illumina adapters for sequencing.

A DNA pool synthesized from Twist Bioscience was PCR amplified by mixing 1 µL of 1 ng/µL of the pool, 1 µL of 10 µM forward primer, 1 µL of 10 µM reverse primer, 10 µL of 2X KAPA HIFI enzyme mix, and 7 µL of molecular grade water. PCR was performed in a thermocycler with the following protocol (1) 95°C for 3 min, (2) 98°C for 20 s, (3) 56°C for 20 s, (4) 72°C for 20 s, (5) go to step 2 for about 15 cycles, (6) 72°C for 30 s. The amplified product was PCR purified using QIAGEN PCR Purification Kit (Cat No: 28104). The sample concentration was measured using Qubit 3.0 fluorometer.

This enriched Twist pool was mixed with 100 times more of the Forward Primer (e.g., [FP]/[pool]=100) at 500 nM of the pool. 20 µL of this mixture was mixed with 20 µL of 2X KAPA HIFI enzyme mix, followed by linear amplification with the following protocol: (1) 95°C for 3 min, (2) 98°C for 20 s, (3) 62°C for 20 s, (4) 72°C for 20 s, (5) go to step 2 for 2 time, and (6) 72°C for 30 s. The mixture contains 250 nM of double-stranded DNA (dsDNA) and less than 750 nM of single-stranded DNA (ssDNA).

The sample was linearly amplified again using an Internal Primer (IP) by mixing 40 µL of the 250 nM dsDNA mixture, 12 µL of 10 µM Internal Primer (IP), and 12 µL of 2X KAPA HIFI enzyme mix. Linear amplification was performed with the following protocol: (1) 95°C for 3 min, (2) 98°C for 20 s, (3) 56°C for 20 s, (4) 72°C for 20 s, and (5) 72°C for 30 s. The mixture contains 156 nM of fully dsDNA pool and less than 468 nM of partially dsDNA with feature region exposed (feature region will hybridize to a query strand).

6.4 uL of the mixture (containing 156 nM of the fully dsDNA pool and 468 nM of partially dsDNA) was mixed with 1 uL of a query strand at 10 nM, 10 uL of 2 M sodium chloride buffer and 2.6 uL molecular grade water. This resulted in an 1:100 ratio of query to the fully dsDNA pool and a final concentration of the fully dsDNA pool at 50 nM. This mixture was annealed in a thermocycler by first heating up to 95°C for 3 mins, and then slowly cooling down to 21°C at the rate of 1°C per 20 min.

3 ug of Streptavidin-coated magnetic beads (Dynabeads MyOne Streptavidin T1, Thermo Fisher Scientific) was used for 1 pmole of a query strand. The beads were washed 3 times in binding and washing buffer (5 mM Tris-HCl (pH 7.5), 0.5 mM EDTA, 1 M NaCl), then added to the hybridization sample at room temperature. After incubating at room temperature for 15 minutes, the samples sat on a magnet rack to recover the beads and binding DNA. The supernatants were removed, and the beads were washed 3 times using 100 µL of binding and washing buffer. The beads binding DNA was resuspended in 50 µL 1X elution buffer containing 10 mM tris-Cl, at pH 8.5. The resuspended samples were digested using USER enzyme by mixing 50 µL of the sample with 2 µL of USER enzyme, and 5.8 µL of NEB 10X cut smart buffer at 37°C for 20 minutes. The sample sat on a magnetic rack for 1 minute, and the supernatants was recovered.

2 µL of the recovered solution from last step was mixed with 1 uL of 10 uM forward primer, 1 µL of 10 µM reverse primer (RP), 10 µL of 2X KAPA HIFI enzyme mix, and 6 µL of molecular grade water for PCR. PCR was performed in a thermocycler with the following protocol (1) 95°C for 3 min, (2) 98°C for 20 s, (3) 62°C for 20 s, (4) 72°C for 20 s, (5) go to step 2 for a varying number of times depending on the recovery yield of beads extraction, and (6) 72°C for 30 s. 2 µL of the amplified sample was mixed with 1 µL of 10 µM forward primer with an overhang of a randomized region (25N), 1 µL of 10 µM reverse primer (RP), 10 µL of 2X KAPA HIFI enzyme mix, and 6 µL of molecular grade, followed by the following thermocycling protocol: (1) 95°C for 3 min, (2) 98°C for 20 s, (3) 62°C for 20 s, (4) 72°C for 20 s, (5) go to step 2 for a varying number of times depending on the recovery yield. This PCR step added a randomized region to the sample for the diversity need of Illumina NextSeq. The size of the PCR product was verified using QIAGEN bioanalyzer. The amplified product was ligated to Illumina sequencing adapters with TruSeq Nano reagents and protocol. The ligated samples were sequenced using Illumina NextSeq.

**Fig. S1.**
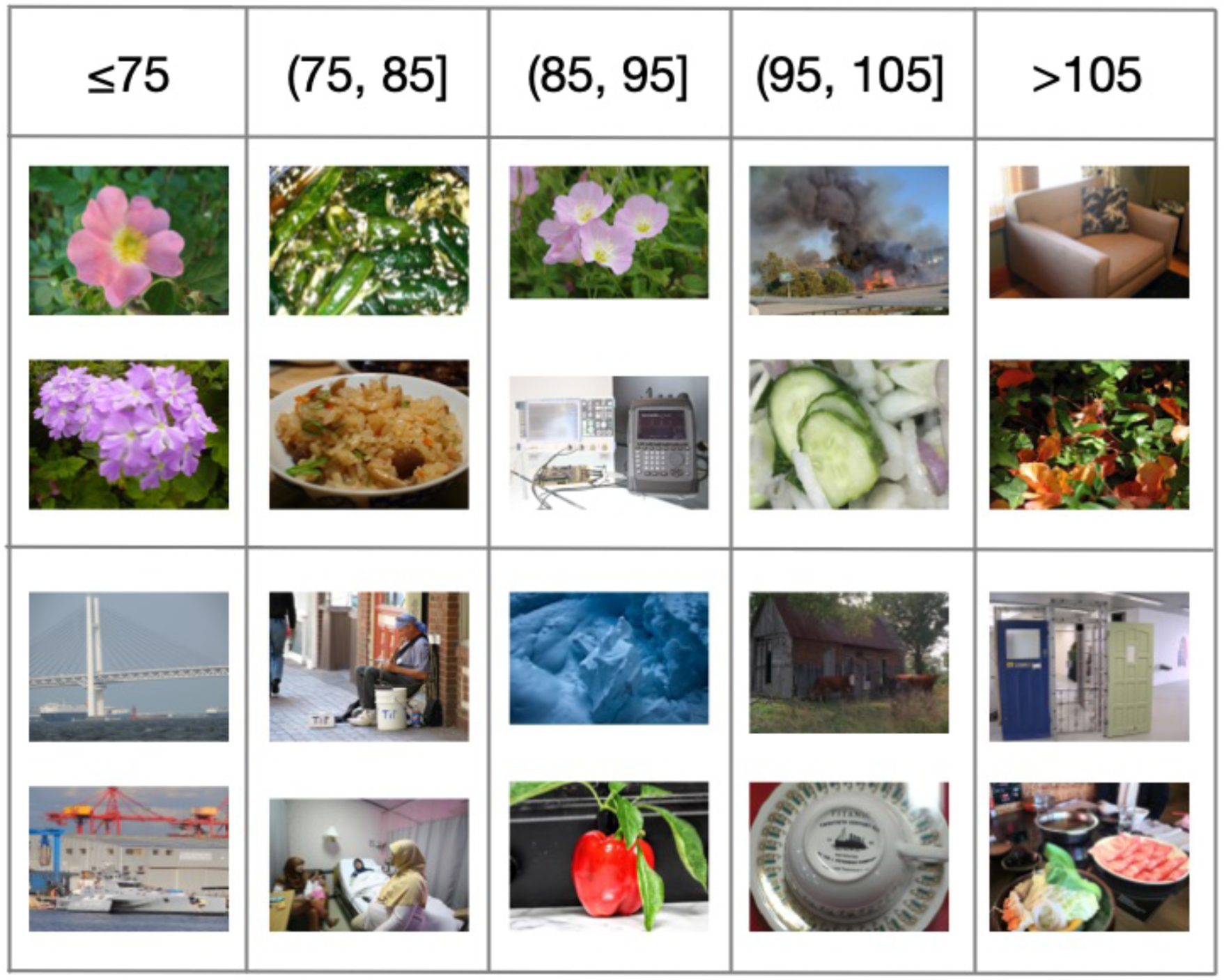
Illustration of the relationship between pairwise feature-vector Euclidean distance and pairwise subjective similarity. Each column represents a range of Euclidean distances, and each row depicts a pair of images where their feature-vector Euclidean distance falls in that range.

**Fig. S2.**
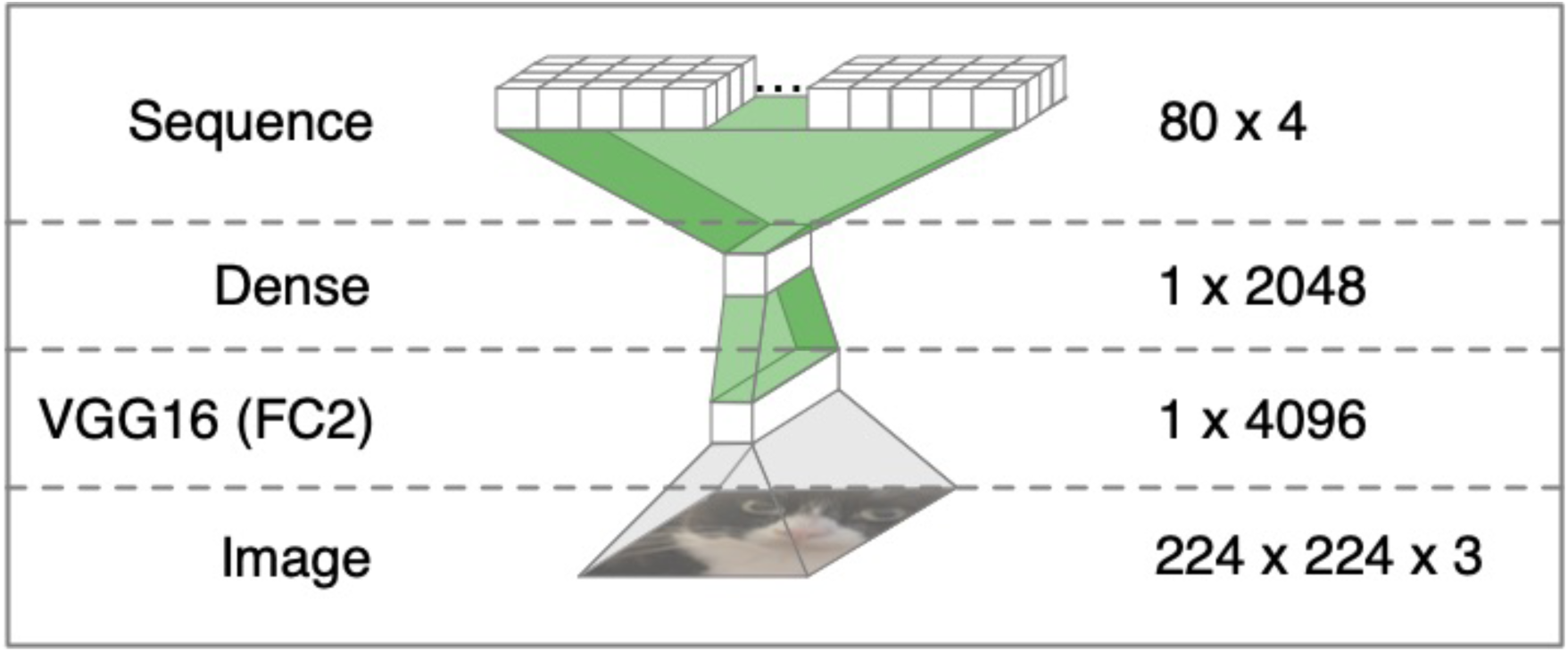
Structure of the sequence encoder network. Layers are opaque, and transformations are translucent. The dimensionality of each layer is shown on the right. Only the transformations highlighted in green have parameters that change during training.

**Fig. S3.**
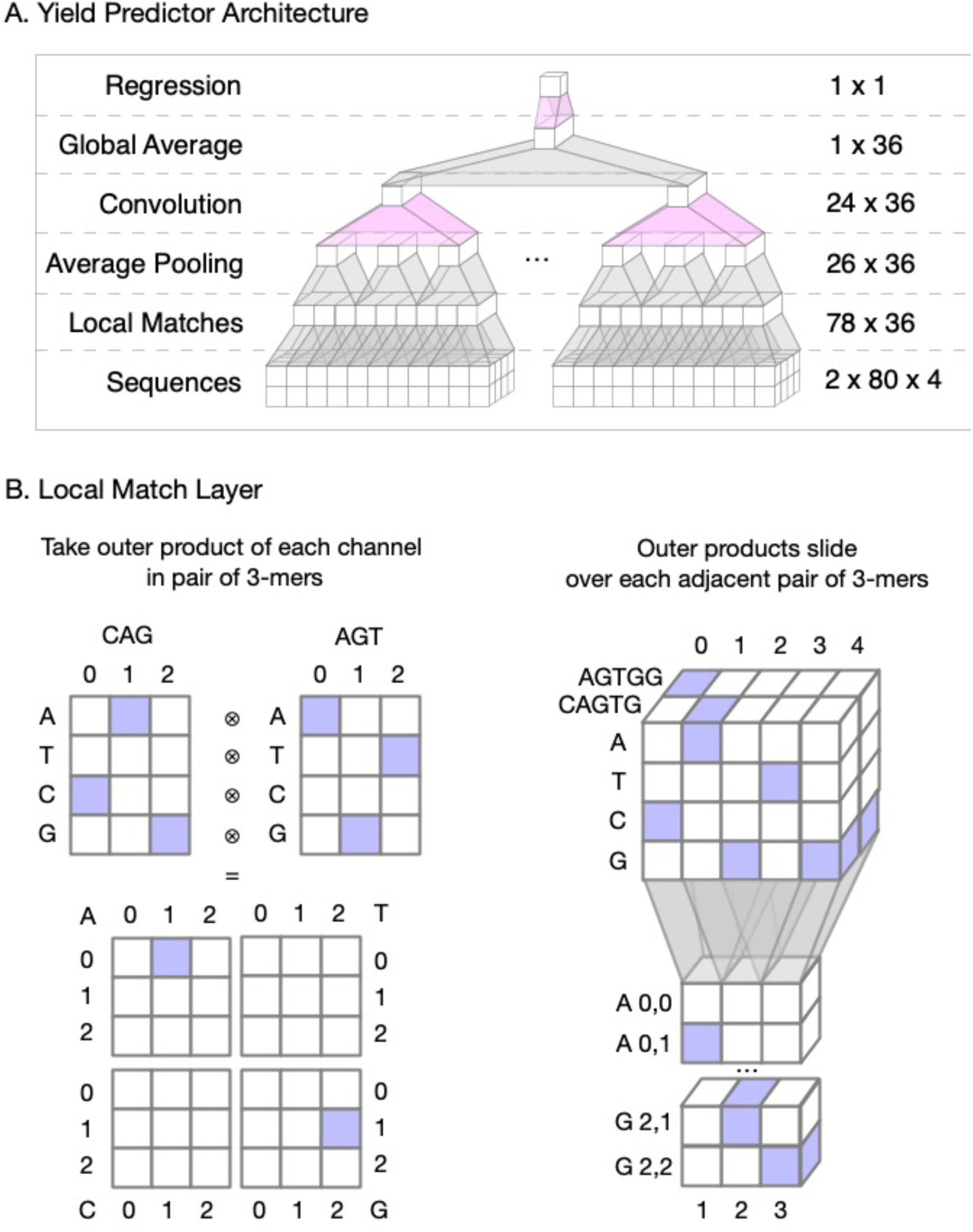
(A) Structure of the yield predictor network. Only the transformations highlighted in pink have parameters that change during training. (B) Illustration of the “local match” operation. Blue cells have a value of 1, and white cells have a value of 0.

**Fig. S4.**
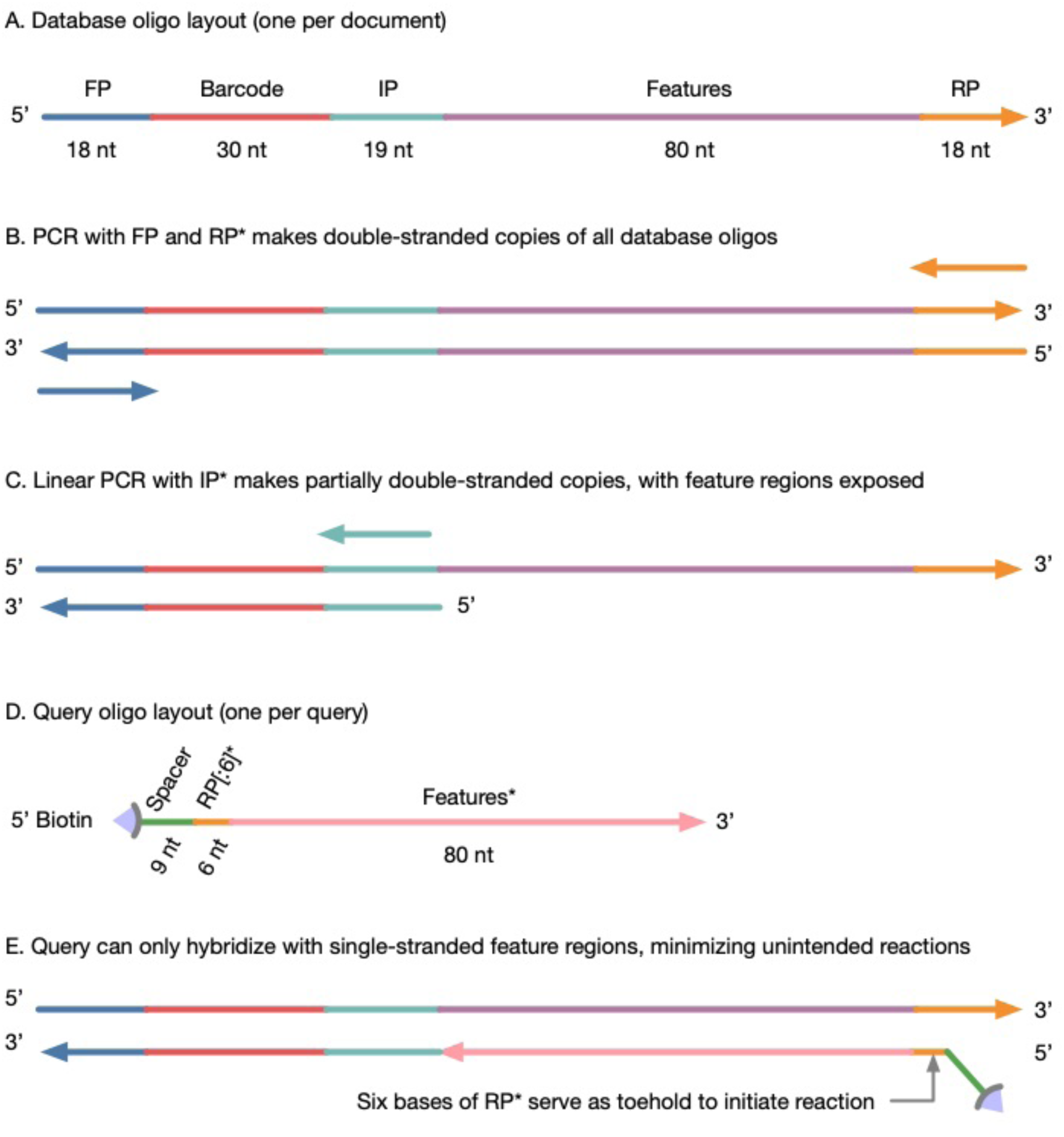
Layouts of single-stranded oligomers and intended double-stranded complexes. Arrowheads indicate 3’ ends of DNA. Asterisks (*) indicate the reverse complement of a DNA sequence.

**Fig. S5.**
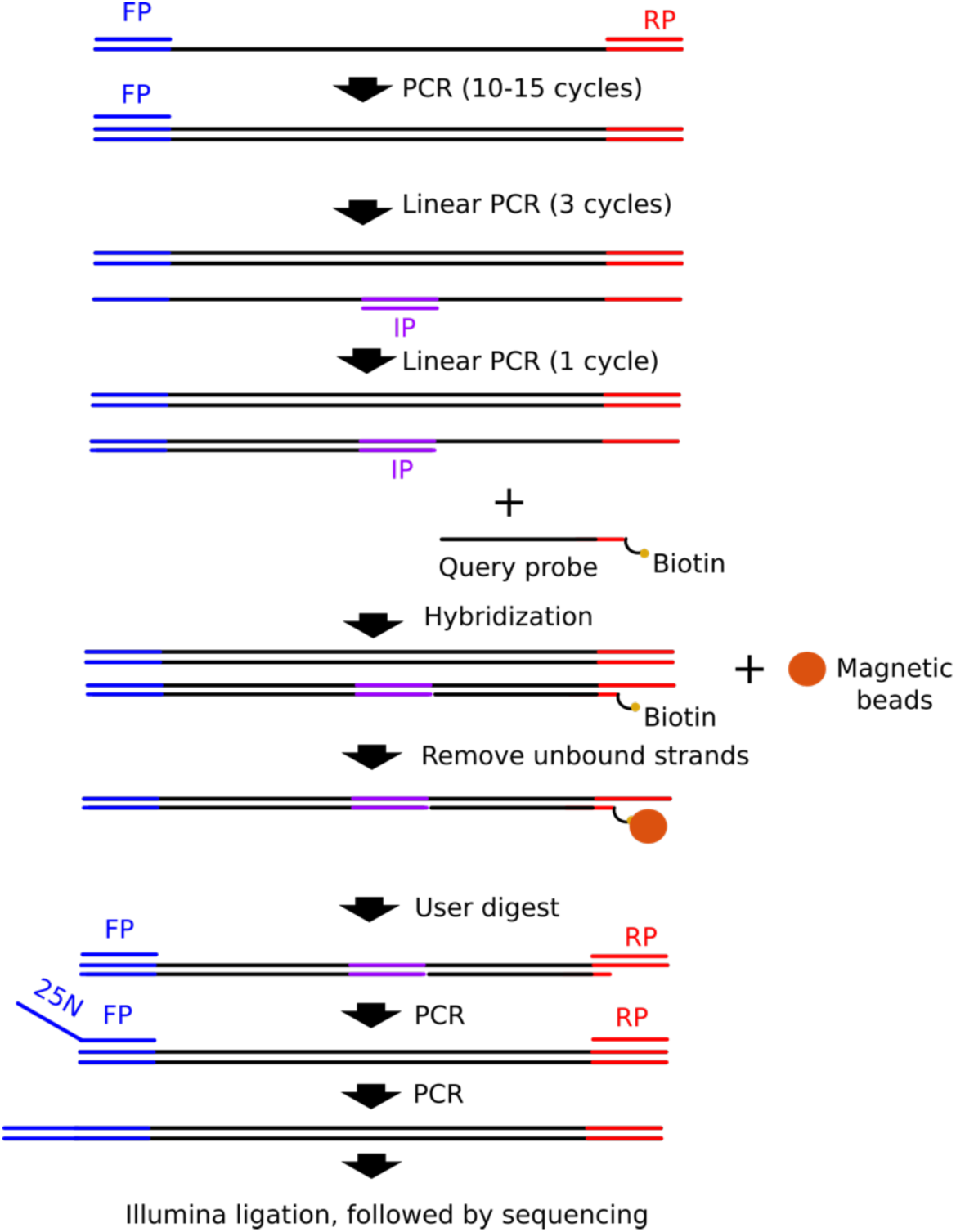
Workflow of a similarity search experiment. A large DNA pool is PCR amplified using a forward primer (FP) and a reverse primer (RP). The enriched product is linearly amplified using the forward primer for 3 cycles. The sample is linearly amplified using an internal primer (IP) to make partially double-stranded copies, with feature region exposed. This mixture is then hybridized with a query strand, followed by magnetic bead extraction. The extracted strands are released from the beads using USER enzyme digestion. The released sample is PCR enriched using FP and RP. The sample is PCR again using RP and FP with a 25N overhang to create a randomized region for the diversity need of Ilumina NextSeq. The sample is ligated to Illumina adapter, followed by next-generation-sequencing.

